# Spinal facilitation of descending motor input

**DOI:** 10.1101/2023.06.30.547229

**Authors:** Giuliano Taccola, Roger Kissane, Stanislav Culaclii, Rosamaria Apicella, Wentai Liu, Parag Gad, Ronaldo M. Ichiyama, Samit Chakrabarty, V. Reggie Edgerton

## Abstract

Highly varying patterns of electrostimulation (Dynamic Stimulation, DS) delivered to the dorsal cord through an epidural array with 18 independent electrodes transiently facilitate corticospinal motor responses, even after spinal injury. To partly unravel how corticospinal input are affected by DS, we introduced a corticospinal platform that allows selective cortical stimulation during the multisite acquisition of cord dorsum potentials (CDPs) and the simultaneous supply of DS. Firstly, the epidural interface was validated by the acquisition of the classical multisite distribution of CDPs on the dorsal cord and their input-output profile elicited by pulses delivered to peripheral nerves. Apart from increased EMGs, DS selectively increased excitability of the spinal interneurons that first process corticospinal input, without changing the magnitude of commands descending from the motor cortex, suggesting a novel correlation between muscle recruitment and components of cortically-evoked CDPs. Finally, DS increases excitability of post-synaptic spinal interneurons at the stimulation site and their responsiveness to any residual supraspinal control, thus supporting the use of electrical neuromodulation whenever the motor output is jeopardized by a weak volitional input, due to a partial disconnection from supraspinal structures and/or neuronal brain dysfunctions.

## Introduction

Dynamic Stimulation (DS) is a highly varying biomimetic protocol of spinal electrical stimulation sampled from an EMG trace recorded from hindlimb muscles during stepping, and delivered to the dorsal spinal cord through an innovative epidural multielectrode array (MEA) provided with low-impedance and fully independent electrodes (Gad et al., 2013; Chang et al., 2014; Taccola et al., 2020a). In anesthetized neurologically intact rats, DS facilitates motor responses evoked by low intensity pulses delivered to both motor cortex and spinal cord, eliciting a significantly higher level of muscle recruitment instead of barely detectable muscle contractions (Taccola et al., 2020a). Furthermore, in spinal cord injured rats acute delivery of DS to spinal segments across the lesion site (Taccola et al., 2020a,b; Culaclii et al., 2021) leads to recovery of motor output. Both effects persist for even a couple of minutes after the end of stimulation, albeit the mechanisms behind these transient modulatory events are still under investigation. Mainly, it must be clarified whether neuromodulation recruits spared, but functionally dormant, fibers at the site of injury, and/or if it improves the responsiveness of spinal interneurons below the lesion, which can respond to other, even weak, inputs that would be otherwise unable to recruit motor pools.

A corticospinal motor command can be dissected out into two main components: a presynaptic input that travels along descending fibers to eventually depolarize the synaptic terminal, and the post-synaptic activation of spinal interneurons. In turn, these spinal neurons will first process the volitional motor input and then pass it on to an ample spinal circuit. Still, the precise identity of these spinal interneurons isn’t clear, although attempts are made to classify them using genetic manipulation of spinal circuitries (Kiehn, 2006) and other experimental approaches (Bellardita et al., 2018; Hsu et al., 2023).

However, besides defining the distinct neuronal chain recruited by corticospinal motor commands, it would be pivotal to clarify how corticospinal input are affected by external interventions, such as electrical spinal neuromodulation. In a perspective to improve the recovery of volitional motor control after SCI, identifying the pre– and/or postsynaptic action sites of neuromodulation could help target the promising DS neurorehabilitative strategy at either spared fiber at levels of injury, or at spinal networks located below the injury site.

Classical electrophysiology approaches can trace the arrival of incoming sensory input to the cord, by dissecting out each different peak composing focal potentials elicited by electrical stimulation and recorded from the surface of the spinal cord. Although the exact profile of focal potentials derived from the cord surface depends on multiple factors, such as recording site, stimulated structures and pulse strength, they always appear as an initial positive spike followed by negative waves reflecting the mono-and polysynaptic activation of spinal neurons followed by a large and long duration positive potential generated by the depolarization of primary afferents (Brooks and Eccles, 1947; Bernhard, 1952, 1953; Eccles et al., 1954; Coombs et al., 1956; Peets et al., 1984; Bozdoğan and Alpsan, 1987; Edgley and Jankowska, 1987; Harrison et al., 1988; Koerber et al., 1990, 1991; Jankowska and Riddell, 1993; Wall and Lidierth, 1997; Riddell and Hadian, 1998; Quiroz-González et al., 2011, 2014; Van Soens et al., 2015). When focal potentials are evoked by electrical pulses applied to peripheral nerves and are recorded from the dorsal horns, they are named cord dorsum potentials (CDPs). Moreover, Jordan and colleagues also described CDPs elicited by electrical stimulation of supraspinal structures (Noga et al., 1995), i.e., the locomotor mesencephalic region, as composed of four positive waves (P_1-4_) and a slow negative wave (N; Noga et al., 1995). In particular, the P_1_ peak has been attributed to the arrival of the descending volley to the cord (pre-synaptic input), followed by subsequent P-waves that are produced by the sequential recruitment of spinal interneurons (post-synaptic activation). In particular, the P_2_ wave corresponds to the monosynaptic activation of spinal interneurons by descending input, while the P_3_ wave corresponds to the di– or tri-synaptic activation of spinal interneurons. The latency for the onset of the P_4_ wave could be caused by some input travelling through slow conducting fibers. However, this is harder to determine since the P_4_ wave follows the extended negative wave N, which likely reflects a presynaptic inhibition of primary afferent fibers. Interestingly, the amplitude of P_2_ to P_4_ waves increases when spinal networks are activated during the appearance of fictive locomotion (Noga et al., 1995). In the present study, we replicated a similar approach while delivering DS, to explore whether neuromodulation mainly facilitates the transit of descending pre-synaptic pulses or, alternatively, increases the excitability of post-synaptic spinal interneurons, which in turn become more responsive to weak spared cortical commands.

Two unique technological features were adopted to achieve this aim: (1) an interface for the punctiform stimulation of the motor cortex to selectively activate the descending volitional motor commands to hindlimb muscles (Russell et al., 2019), and (2) an epidural MEA with independent electrodes able to stimulate the cord with DS (Gad et al., 2013; Chang et al., 2014; Taccola et al., 2020a) that here, for the first time, we adopted also to record CDPs from multiple sites of the adjacent dorsal cord. We first validated our technology for the simultaneous recording of multiple CDPs in response to electrical stimulation of peripheral nerves, similarly to what already reported using glass or metal ball single electrodes (Brooks and Eccles, 1947; Bernhard, 1952, 1953; Coombs et al., 1956). Then, we introduced a proof of concept for an innovative corticospinal platform, which is composed of two elements. The first is a cortical stimulating interface (Russell et al., 2019) for the selective contralateral muscle recruitment of hindlimb muscles following punctiform delivery of electrical pulses to the motor cortex. The second element of the corticospinal platform is the simultaneous acquisition of multisite CDPs through the same epidural MEA that is used to supply DS. To this aim, we continuously delivered descending test pulses before and after applying DS at sub-threshold intensities to the cord. Analysis of CDP peaks identified a selective post-synaptic facilitation of the short latency P_2_ component, without any changes in descending volleys (P_1_). This phenomenon suggests that the main mechanism of neuromodulation for the recovery of volitional motor control after SCI relies on a subthreshold excitation that both increases the excitability of sensorimotor spinal networks located at the stimulation site and their responsiveness to any residual supraspinal control, which adds on to the well-known enhanced integration of sensory inputs (Courtine et al., 2009). Once motor threshold is reached, more motor units are eventually recruited even by weak spared input passing through the lesion (Gerasimenko et al., 2015). This implies that remaining descending connections to neuronal networks located below the lesion persist in a significant number of individuals with complete, chronic paralysis (Harkema et al., 2011), in accordance with findings in postmortem histological analysis (Kakulas and Kaelan, 2015).

## Materials and methods

The present work explores the focal responses arising on the surface of the dorsal spinal cord following selective cortical stimulation. High varying patterns of current (Dynamic Stimulation, DS) sampled from EMGs during locomotion were delivered to spinal networks via an ad hoc epidural electrode array (Gad et al., 2013, Chang et al., 2014). Briefly, we recorded the topographic distribution and input-output profile of focal potentials recorded from the dorsal surface of the spinal cord in response to electrical stimulation of two distinct peripheral nerves, to verify whether our interface reproduced the classical volleys of cord dorsum potentials (CDPs). We subsequently explored the dynamics of the multipeak CDPs and muscle EMGs during corticospinal input before and after DS, uncovering novel correlations between the volley profile of CDPs and muscle recruitment.

### Experimental design

Data were collected from 24 adult male and female Wistar rats (250 – 300 g body weight) anesthetized by intraperitoneal (IP) administration of a ketamine (100mg/kg) and xylazine (5mg/kg) mix. At the end of experiments, animals were euthanized with either isoflurane or sodium pentobarbital (IP, 80-100 mg/Kg) followed by cervical dislocation.

All procedures were approved by the UK Home Office and performed under the Animals (Scientific Procedures) Act 1986 and by the Animal Research Committee at UCLA in accordance with the guidelines of the National Institutes of Health (NIH) Guide for the Care and Use of Laboratory Animals and with the European Union directive on animal experimentation (2010/63/EU).

### Intramuscular EMG electrode implantation and recordings

Animals were kept under anesthesia over a heating pad (37° C) during the entire duration of surgery and experiments. EMG responses were recorded from left and right tibialis anterior (TA) muscles using wire electrodes (40 AWG; 79 μm diameter) insulated with polyurethane. Electrodes were mechanically stripped of insulation, 1mm from the tips, and inserted through the shank skin using a 23 G needle running the wires through the mid-belly of the muscle. A common ground was inserted subcutaneously in the mid-back region for EMGs. Signals were acquired using a differential amplifier (D160®, Digitimer Ltd, Hertfordshire, UK; Gain x2000, filtered 10-1000 Hz), notched (Hum Bug®, A-M Systems, Sequim, WA, US), digitalized at 20, 30 or 40 KHz (Micro1401-3®, Cambridge Electronic Design Limited, Cambridge, UK) and stored in a PC for following off-line analysis (Signal® version 5.09, Cambridge Electronic Design Limited, Cambridge, UK).

### Epidural multi electrode array implantation

To simultaneously deliver patterns of intrinsically varying signals to multiple segments of the spinal cord we used a high-density platinum-based multi-electrode array consisting of three longitudinal columns and six horizontal rows of independent low-impedance electrodes (total of 18 independent electrodes; Gad et al., 2013; Cheng et al., 2014; Taccola et al., 2020a). The high-definition and spinal precision of the epidural interface was demonstrated by the selective activation of extensor or flexor motor pools while varying bipolar stimulation parameters (Taccola et al., 2020a).

To implant the multi-electrode array in the dorsal epidural space, a T12 to L2 vertebrae laminectomy was performed to expose the spinal cord. After epidural placement, the array was covered with small cotton balls rinsed in saline. Back muscles and skin were sutured using 5.0 Vicryl® (Ethicon, New Brunswick, NJ, USA), with leads to the array exiting through the skin. A common ground for all array electrodes, independent from EMG ground, was inserted subcutaneously in the left forearm. Throughout the text, segmental levels of the spinal cord are indicated in accordance to the topography of spinal roots entrance, as previously reported (Lavrov et al., 2008).

### Electrical stimulation protocols

Our electrical stimulation approach closely followed the protocol in our previous work (Taccola et al., 2020a). In summary, our stimulation waveforms reused the Dynamic Stimulation (DS) signal, which was previously created from an EMG recording from the Sol muscle of a neurologically intact adult rat walking on a treadmill. The recording was duplicated into two channels which are time-staggered, and then digitized into an ASCII text file, and modified to be compatible with our programmable stimulation device (STG® 4008; Multi Channel Systems, Reutlingen, Germany) and its associated proprietary software (MC Stimulus II). As explained in our past reports, the DS protocol is an electrically safe charge-balanced stimulation, ensured by the method by which the protocol was created. The two-channel, time-staggered stimulus protocol was simultaneously delivered through two outputs to the left and right external columns of array electrodes with opposite rostro-caudal cathode location. At each experiment, the rostral cathode was indifferently placed on the right or on the left columns of the array and then a rostral anode was consequently placed on the other side. DS protocol was applied at four peak-to-peak amplitudes: 400, 500, 600, 800 µA, which were adjusted by the stimulator device’s software.

### Peripheral nerve stimulation

For plantar nerve electric stimulation, a pair of needle electrodes were placed subcutaneously at the ankle behind the medial malleolus and 1 cm proximal. The peroneal nerve was stimulated distally at the ankle and proximally at the popliteal fossa, using a cuff electrode. Single biphasic stimuli (duration 100 – 200 µs) were supplied at 0.3 Hz (808 ± 89 µA, ISO-STIM-01M®, npi electronic GmbH, Tamm, Germany).

### Electrical stimulation of motor cortex

Under stereotaxic coordinates a 5 x 5 mm craniotomy site expose the motor cortex, and unilateral cortical stimulation was performed by implanting an epidural cortical array (Russel et al., 2018). Using our custom array (Russel et al., 2018) an epoch of three or five pulses (inter stimulus interval, ISI = 3 ms) were supplied at 0.3 Hz (808 ± 89 µA, ISO-STIM-01M®, npi electronic GmbH, Tamm, Germany) were applied to the hindlimb region of the motor cortex (Neafsey et al., 1986). Forelimb and shoulder muscles were monitored to ensure no spillover effects arising from cortical stimulation during targeted hindlimb stimulation. Similarly, the absence of ipsilateral TA contractions was confirmed by continuous EMG recordings.

### Data analyses

The amplitudes and the time to peak of each CDP volley were individually calculated. To plot time course graphs, the amplitude of each P2 peak was determined for consecutive sweeps. For statistical comparison, 30 peaks were averaged immediately before and after the application of DS.

### Statistical Analysis

Data are indicated as mean ± SD values, with *n* referring to the number of experiments. After determining the normality of the distribution of data based on a Kolmogorov-Smirnov normality test, statistical analysis was performed using SigmaStat® 3.5 software (Systat Software, San Jose, CA, USA) to compare the mean ± SD of different experimental conditions. In the current study, all values collected from each experiment before and after DS were parametric and then data were analyzed using Student’s paired t-test. Results reached significance when P < 0.05.

## Results

### Peripherally evoked surface potentials are recorded from the dorsal cord

We explored the possibility to simultaneously record electrical potentials from multiple sites of the lumbar enlargement through electrodes of the epidural MEA, which have to date only been used to deliver pulses to the spinal networks (Taccola et al., 2020a,b; 2021; 2022). Thus, terminal experiments were performed in intact adult rats, by stimulating the peroneal and/or plantar nerves on both sides (Kurokawa et al., 2004). In accordance with Bernhard (1952, 1953), the anatomy of a prototypical average CDP is reported in Figure 1A, where a multipeak average volley was derived from the epidural surface of the cord at the midline of spinal level L6, in response to single square pulses (duration = 0.2 s) delivered to the left peroneal nerve. Upon each pulse delivery, a biphasic artifact of stimulation occurs, followed by a downward deflection after 1.8 ms, which has been attributed to the afferent volley (AV) entering the spinal cord via lumbar dorsal roots (DRs). The following positive trough corresponds to the depolarization of presynaptic terminals occurring 2.3 ms after the stimulus. Then, the baseline rises to reach a first negative focal potential (N1, represented by convention as an upward deflection) that corresponds to the post-synaptic depolarization of spinal neurons by fast conducting group IA fibers (Coombs et al., 1956). According to classical spinal neurophysiology, the latency between presynaptic depolarization and the onset of the subsequent N1 peak corresponds to the synaptic delay, which is almost 0.3 ms in our sample averaged trace, not dissimilar from the 0.4 – 0.5 ms originally calculated in cat spinal cords following electrical stimulation of the quadriceps nerve (Brooks and Eccles, 1947; Eccles et al., 1954). After N1, a second delayed negative peak generates a slower and higher potential, known as N2, peaking in our experiments approximately at 2.7 ms, which reflects both, the response to impulses travelling through group II fibers (Coombs et al., 1956), and the di– and tri-synaptic recruitment of a broader population of spinal interneurons. Interestingly, a latency of about 3 ms has been reported for the negative wave in response to the activation of L7 spinal neurons in anesthetized cats through stimulation of group II afferents in the sural nerve (Coombs et al., 1956). While AV and PV describe presynaptic events confined to afferent fibers, N1 and N2 are attributed to the postsynaptic activation of spinal neurons. The CDP volley elicited by peripheral electrical stimulation terminates with a slow positive P wave, attributed to inhibitory input that follow the primary afferent depolarization of close DRs (Quiroz-González et al., 2011; 2014). The main contributions to each phase of the CDP evoked by peripheral electrical stimulation are summarized in Figure 1B.

**FIGURE 1.**
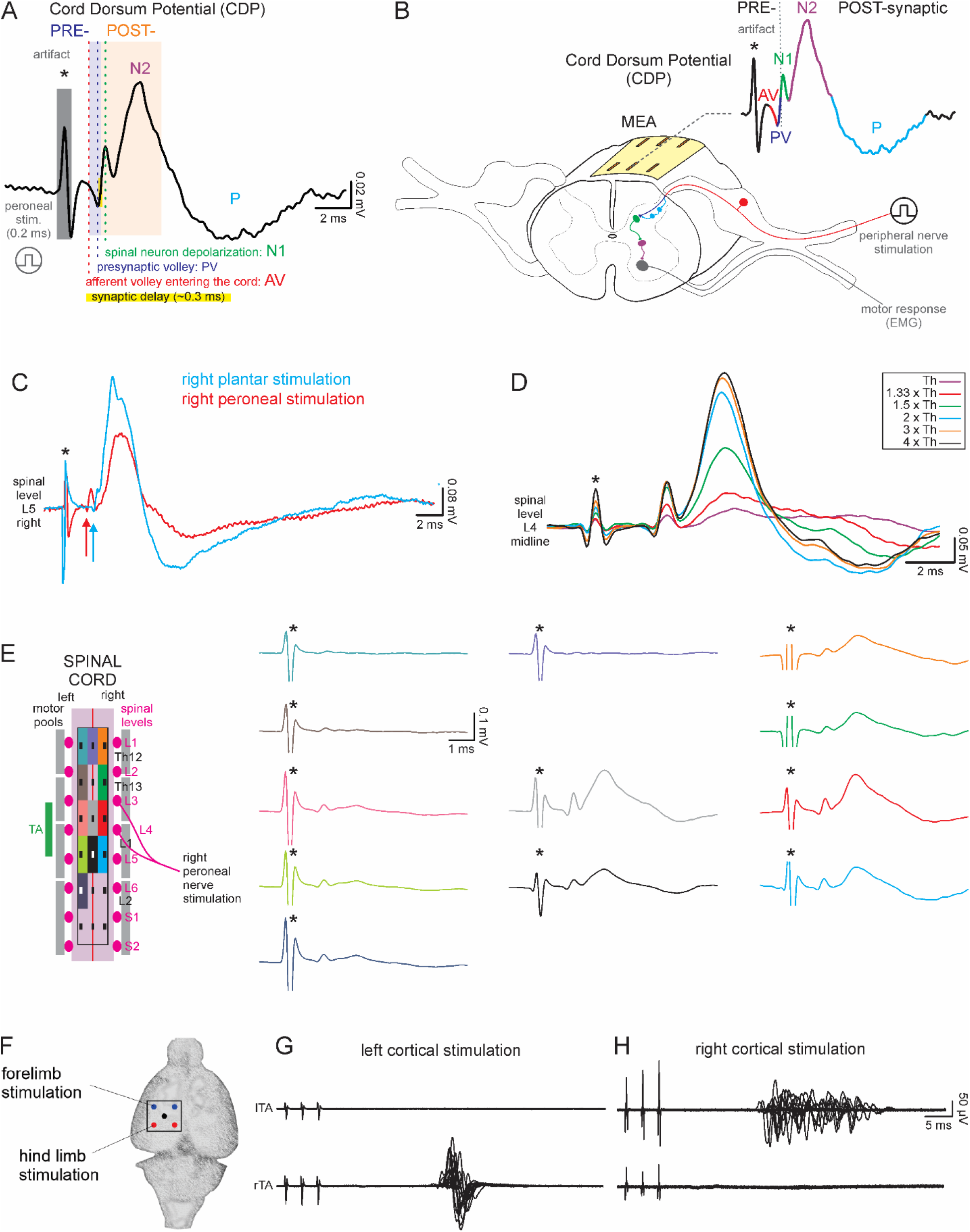
Two distinct epidural multi-electrode arrays were adopted for both recording cord dorsum potentials elicited by peripheral nerve stimulation and selectively stimulating the motor cortex. **A**. An average CDP profile recorded from spinal level L6 at midline and induced by square pulses (intensity = 0.03 V; duration = 0.2 s) delivered to the left peroneal nerve is conventionally reported with negative potentials pointing upwards. Following a biphasic artifact of stimulation (grey field), signal is divided in two fields, i.e. pre–(pale purple) and post-synaptic spinal response (pale orange). Main CDP components are indicated by conventional letters (AV, PV, N1, N2, P). The latency between presynaptic depolarization and onset of the subsequent N1 peak reflects the synaptic delay, which is here close to 0.3 ms. **B**. The cartoon shows a segment of the spinal cord with the epidural array placed on the surface of the dorsal cord. Each peak of the CDP is displayed according to the same color code used in the adjacent schematized spinal circuitry recruited by afferent stimulation. **C**. Two averaged CDPs collected from the right L5 spinal segment of the same animal are superimposed in response to electrical pulses serially delivered to the right plantar nerve (cyan trace, intensity = 0.04 V, duration = 0.2 ms) and the right peroneal nerve (red trace, intensity = 0.1 V, duration = 0.2 ms). Colored arrows indicate the shortest latency of the CDP elicited by peroneal pulses. **D**. Superimposed mean traces acquired from the midline of the L4 spinal segment in response to input/output stimulation of the left peroneal nerve (0.3 Hz, duration = 0.2 ms). Intensity of stimulation is reported in the box as times x threshold (Th = 0.015 V). **E**. A schematic view of the multielectrode array placed on the dorsal spinal cord. Dimensions of the array and electrodes are calibrated on the width and length of the cord, as well as on the root entry positions. In response to right peroneal nerve stimulation (*, intensity = 0.06 V, duration = 0.2 ms), simultaneous CDPs from 12 independent electrodes of the array are traced using the same color code as in A. The empty spots correspond to non-working electrodes. Traces in A, C, D and E come from different experiments. **F**. A schematized view of the epidural multielectrode (5 leads) array adopted for selective cortical stimulation. **G**. Simultaneous EMG recordings from left (top) and right (down) tibialis anterior muscles (TAs) in response to a short train with three biphasic pulses (333 Hz, intensity = 55 µA) delivered every 3 s to the left motor cortex. Responses appear only on the contralateral muscle. **H**. By moving the array to the right side of the motor cortex, the train of pulses (intensity = 90 µA) elicits motor responses only on the left TA without any signals from the ipsilateral muscle.

At higher intensities of stimulation, it has been reported the appearance of both, a small negative N3 wave superimposed on the large positive P wave, which might originate from the activation of group III fibers (Bozdoğan and Alpsan, 1987), and of a slower late component (LC) with a latency in the range of 200 ms, which has been ascribed to electrical stimulation of C-fibers in the tail (Peets et al., 1984).

The peak composition of acquired CDPs is quite consistent among multiple rats, with only minor changes in latency corresponding to stimulation of different nerves. Indeed, in the same animal, peroneal and plantar right nerves were serially stimulated while CDPs were derived from the right side of L5 spinal segment (Fig. 1C). In the superimposed average traces (Fig. 1C), the CDP evoked by plantar stimulation (cyan) appears with a more delayed onset (0.5 ms) than that (red) elicited by peroneal pulses (as indicated by staggered arrows centered on PVs), reminiscent of the longer distance and smaller diameter of plantar fibers compared to the peroneal nerve stimulated in the popliteal fossa.

When the amplitude of monopolar pulses applied to the left peroneal nerve increased (duration = 0.2 ms; intensities from 0.015 V, Th, to 0.06 V, 4 x Th), the amplitude of CDP peaks recorded from the midline at L4 spinal level, progressively augmented, while their latencies remained unaffected (Fig. 1D). Notably, for stimulation intensities higher than 1.5 x Th, N2 becomes higher than N1, reflecting the polysynaptic recruitment of a wider interneuronal network at strong intensities of stimulation. The P wave appears for intensities over Th, progressively increasing for stronger stimuli and reaching larger values for 2 x Th pulses, without any further increments at even greater intensities of stimulation. A similar experiment was replicated in 5 rats and results are reported in Table 1 (Th = 11 ± 3 µV).

**Table 1.**
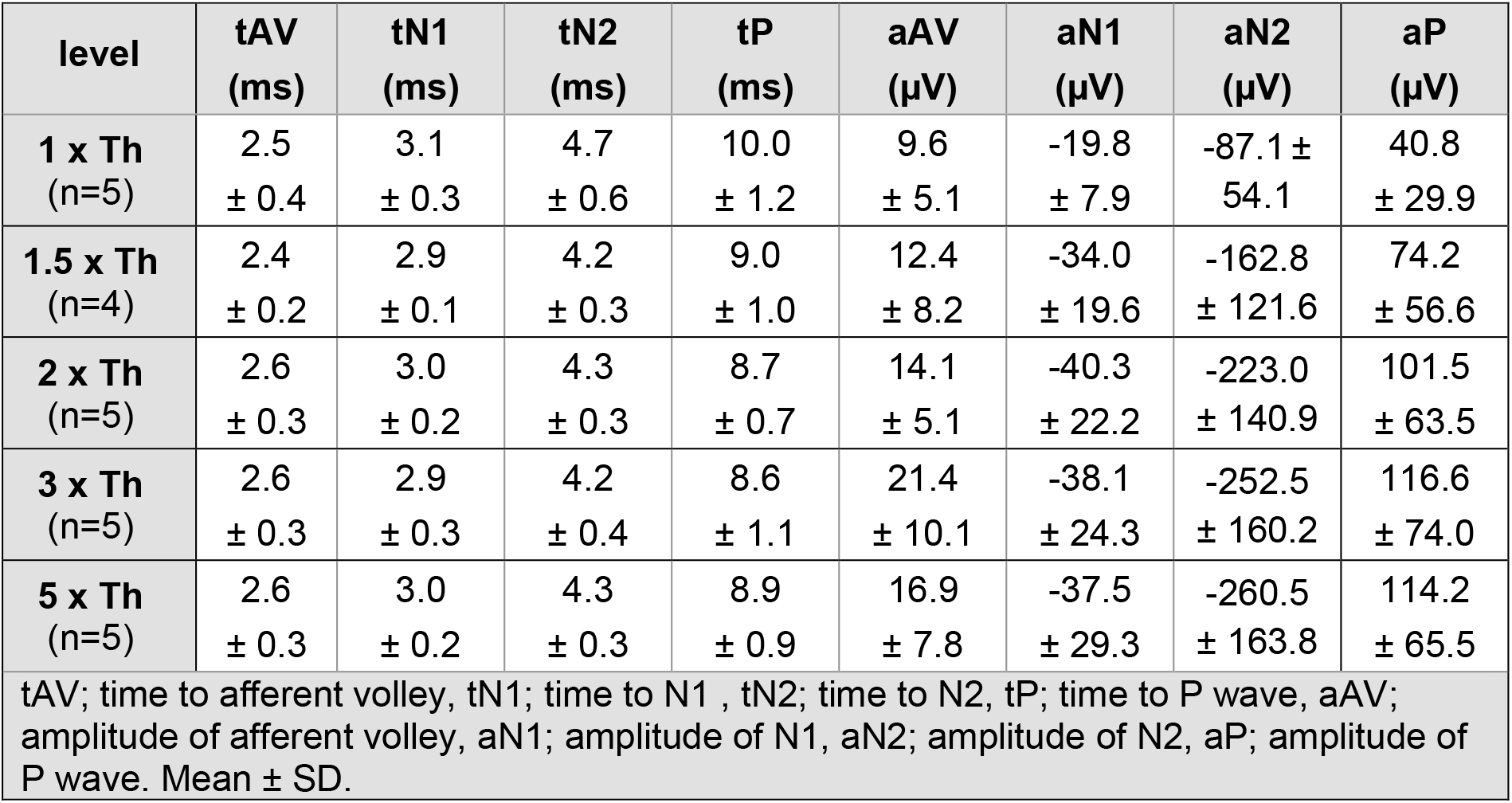
Latency (t) and amplitude (a) of CDP waves elicited through plantar nerve stimulation using pulses of increasing intensities, defined as times of motor threshold.

The MEA adopted in the current study relies on 18 low-impedance electrodes that can be used for multisite stimulation of the cord and for the simultaneous acquisition of potentials from multiple loci. We exploited this unique feature of the interface to map the spinal projections of stimulated nerves. The rostrocaudal location of peak CDP amplitude was quantified to define the highest density of afferent’s boutons (Koerber et al., 1990). In a sample experiment, in response to pulses applied to the right peroneal nerve, CDPs were simultaneously recorded from the twelve sites (Fig. 1E). Averaged traces show only small N1 peaks on the contralateral side of the cord, while both N1 and N2 appear on the right column of the MEA with the highest responses taken from the L3/L4 spinal segments (red trace), according to the topography of spinal targets of peroneal afferents, described through anatomical tracing (Nicolopoulos-Stournaras and Iles., 1983; Koerber et al., 1990). However, at L3/L4 spinal levels, the largest CDP is derived from the electrode placed on the midline (grey trace). This confirms the known midline location of afferent synapses (Bozdoğan and Alpsan, 1987; Smith and Bennett, 1987), also corroborated by the identification of projection patterns of primary afferent fibers that migrate to the midline as they course away from their entry into the spinal cord (Koerber et al., 1990).

For each peak of peripherally evoked CDPs, we measured latency and amplitude values from multiple spinal levels of the lumbosacral cord, homolaterally to the side of stimulation. Pooled data from many experiments are reported in Table 2 and Table 3 for peroneal and plantar stimulations, respectively.

**Table 2.**
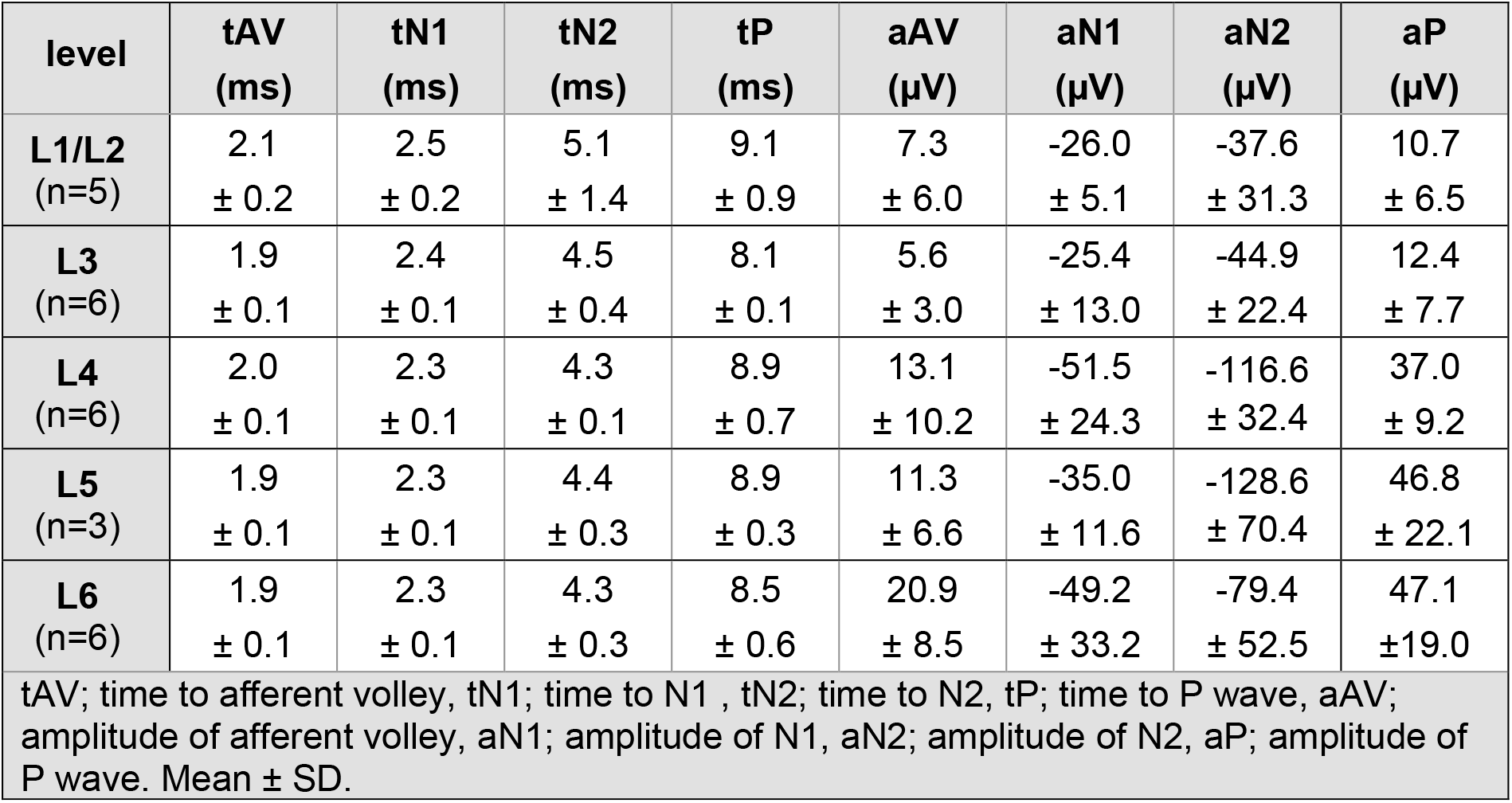
Latency (t) and amplitude (a) of CDP waves recorded from lumbosacral spinal levels and elicited ipsilaterally from the side of peroneal nerve stimulation.

**Table 3.** Latency (t) and amplitude (a) of CDP waves recorded from lumbosacral spinal levels and elicited ipsilaterally from the side of plantar nerve stimulation.

| level | tAV<br>(ms) | tN1<br>(ms) | tN2<br>(ms) | tP<br>(ms) | aAV<br>(μV) | aN1<br>(μV) | aN2<br>(μV) | aP<br>(μV) |
| --- | --- | --- | --- | --- | --- | --- | --- | --- |
| **L1/L2**<br>(n=5) | 2.6<br>± 0.4 | 3.0<br>± 0.3 | 4.5<br>± 0.7 | 8.2<br>± 0.4 | 12.1<br>± 6.0 | -15.1<br>± 7.3 | -61.6<br>± 35.5 | 33.1<br>± 15.8 |
| **L3**<br>(n=9) | 2.3<br>± 0.1 | 2.8<br>± 0.1 | 4.0<br>± 0.2 | 9.6<br>± 0.7 | 13.5<br>± 3.1 | -9.5<br>± 8.3 | -72.0<br>± 16.9 | 33.3<br>± 12.8 |
| **L4**<br>(n=11) | 2.7<br>± 0.3 | 3.0<br>± 0.3 | 4.4<br>± 0.4 | 9.0<br>± 0.7 | 12.1<br>± 9.3 | -19.6<br>± 10.3 | -124.1<br>± 78.1 | 54.2<br>± 28.2 |
| **L5**<br>(n=11) | 2.5<br>± 0.3 | 2.9<br>± 0.3 | 4.3<br>± 0.4 | 8.8<br>± 0.9 | 16.5<br>± 8.9 | -18.3<br>± 13.7 | -139.3<br>± 115.5 | 67.8<br>± 49.0 |
| **L6**<br>(n=3) | 2.5<br>± 0.3 | 2.8<br>± 0.4 | 4.2<br>± 0.4 | 8.5<br>± 1.1 | 14.5<br>± 8.1 | -20.2<br>± 13.1 | -136.2<br>± 95.8 | 60.1<br>± 37.8 |
| **S1/S2**<br>(n=5) | 2.7<br>± 0.2 | 3.1<br>± 0.3 | 4.3<br>± 0.7 | 8.6<br>± 1.3 | 14.4<br>± 0.8 | -12.7<br>± 9.0 | -114.3<br>± 107.6 | 57.9<br>± 29.6 |

tAV; time to afferent volley, tN1; time to N1 , tN2; time to N2, tP; time to P wave, aAV; amplitude of afferent volley, aN1; amplitude of N1, aN2; amplitude of N2, aP; amplitude of P wave. Mean ± SD.

In different animals, segmental CDPs were simultaneously recorded from electrodes located in the same row, and pooled from responses taken from the ipsilateral, middle and contralateral sides of two stimulated peripheral nerves, i.e. peroneal (Table 4; n = 3) and plantar (Table 5; n = 6).

**Table 4.** Segmental CDPs simultaneously recorded from the ipsilateral, middle and contralateral sides of the cord and elicited by peroneal nerve stimulation (n = 3).

| **peroneal nerve stimulation** | **ipsilateral** | **middle** | **Contralateral** |
| --- | --- | --- | --- |
| **tAV (ms)** | 2.0 ± 0.2 | 2.1 ± 0.1 | 1.9 ± 0.1 |
| **tN1 (ms)** | 2.3 ± 0.2 | 2.4 ± 0.1 | 2.4 ± 0.1 |
| **tN2 (ms)** | 4.5 ± 0.1 | 4.2 ± 0.1 | 3.8 ± 0.6 |
| **tP (ms)** | 9.9 ± 0.8 | 9.2 ± 0.1 | 8.9 ± 1.8 |
| **aAV (μV)** | 12.2 ± 9.5 | 23.3 ± 21.0 | 9.4 ± 10.2 |
| **aN1 (μV)** | -45.2 ± 20.1 | -58.1 ± 56.7 | -24.3 ± 15.4 |
| **aN2 (μV)** | -103.7 ± 25.7 | -128.8 ± 141.9 | -30.0 ± 14.0 |
| **aP (μV)** | 36.4 ± 2.2 | 48.99 ± 55.8 | 11.06 ± 5.9 |
| tAV; time to afferent volley, tN1; time to N1 , tN2; time to N2, tP; time to P wave, aAV; amplitude of afferent volley, aN1; amplitude of N1, aN2; amplitude of N2, aP; amplitude of P wave. Mean ± SD. |  |  |  |

**Table 5.**
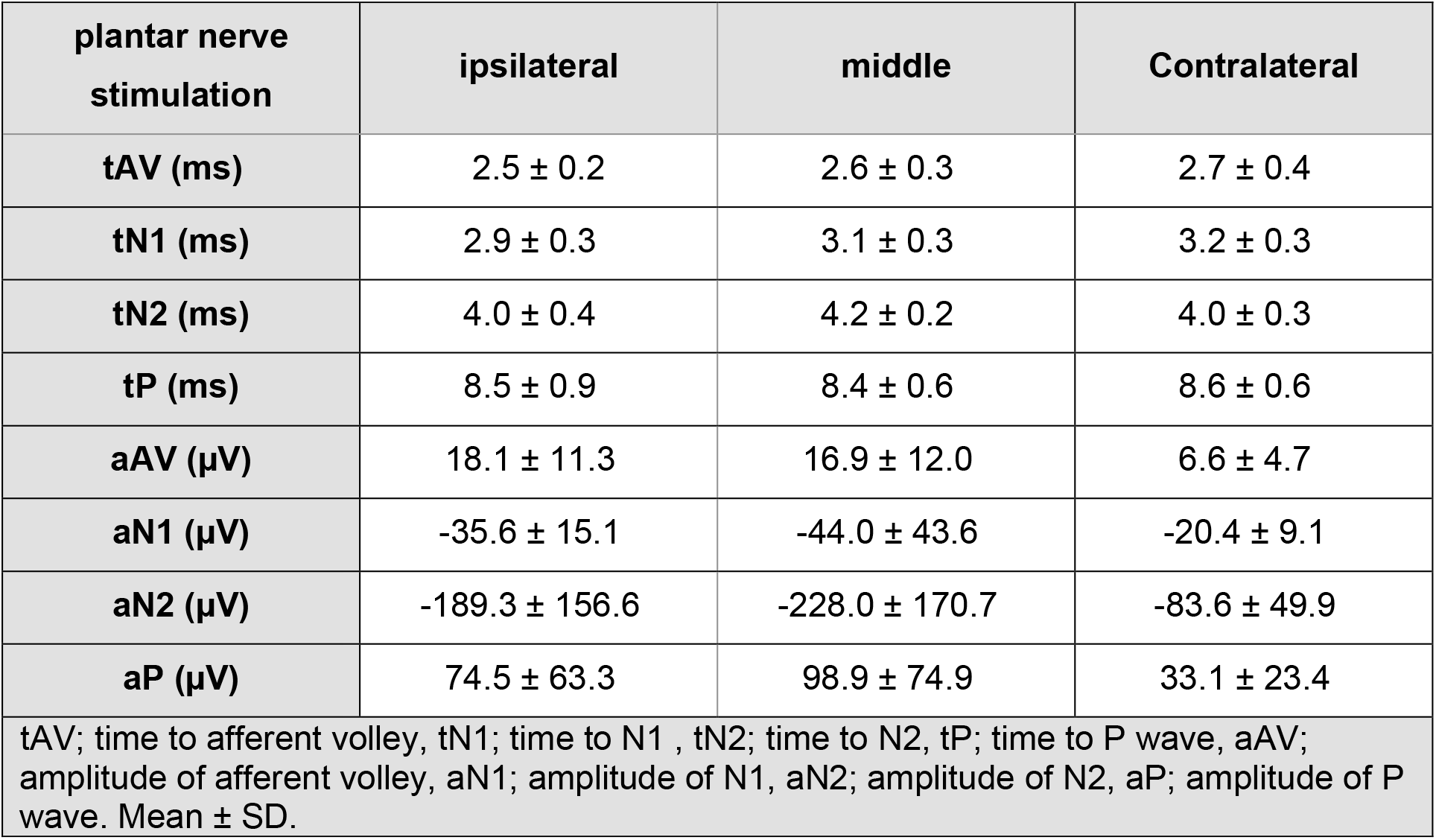
Segmental CDPs simultaneously recorded from the ipsilateral, middle and contralateral sides of the cord and elicited by plantar nerve stimulation (n = 6).

### The selective activation of cortical motor areas through an innovative corticospinal interface evokes simultaneous dorsal volleys from multiple sites of the cord

An innovative array composed of five electrodes (Fig. 1F) was adopted for selective epidural cortical stimulation. Selectivity of the interface was demonstrated by the lack of shoulder contractions in forelimbs (data not shown) and the distinct contralateral activation of TAs when the stimulating MEA was serially placed over leg motor areas (Fig. 1G, left cortical stimulation; Fig. 1H, right cortical stimulation). This observation was replicated in five rats.

To address the action site used by neuromodulation to facilitate volitional motor control, we derived CDPs from the lumbar enlargement in response to descending input elicited by the selective stimulation of hindlimb cortical motor areas. To this aim, we combined the selective cortical stimulating interface (Fig. 2A) and the spinal MEA (Fig. 2B) in an integrated corticospinal platform. As a result, each electrode of the array independently derived superficial focal potentials from the dorsal spinal cord in response to stimulation of a lateral motor area. During selective stimulation of the left motor cortex with brief trains (333 Hz) of three electrical biphasic pulses (duration = 0.2 s), the right TA was serially activated as indicated in the superimposed EMG traces with a latency of 18 – 24 ms from the artifact of the last stimulus (Fig. 2C). When the same experiment was replicated in nine animals, the mean latency of EMG signal was 18.4 ± 6.0 ms. In an exemplar experiment, 16 ms before the appearance of EMG responses, multi peaked CDPs simultaneously occurred in ten sites, spanning across the entire lumbosacral cord (Fig. 2B). Data pooled from six animals reports that the onset of CDPs evoked by corticospinal pulses have a mean latency of 6.0 ± 3.9 ms. When average cortically evoked CDPs are visualized at a higher time-based scale (Fig. 2D), the events recorded along the cord display a stereotyped time-to-peak, but of a different amplitude, with the highest peaks on the right side of the cord. All CDPs have a common structure made of four short-latency positive waves, and a slower and larger negative wave, depicted as superimposed traces from different cord sites (Fig. 2F).

**Figure 2.**
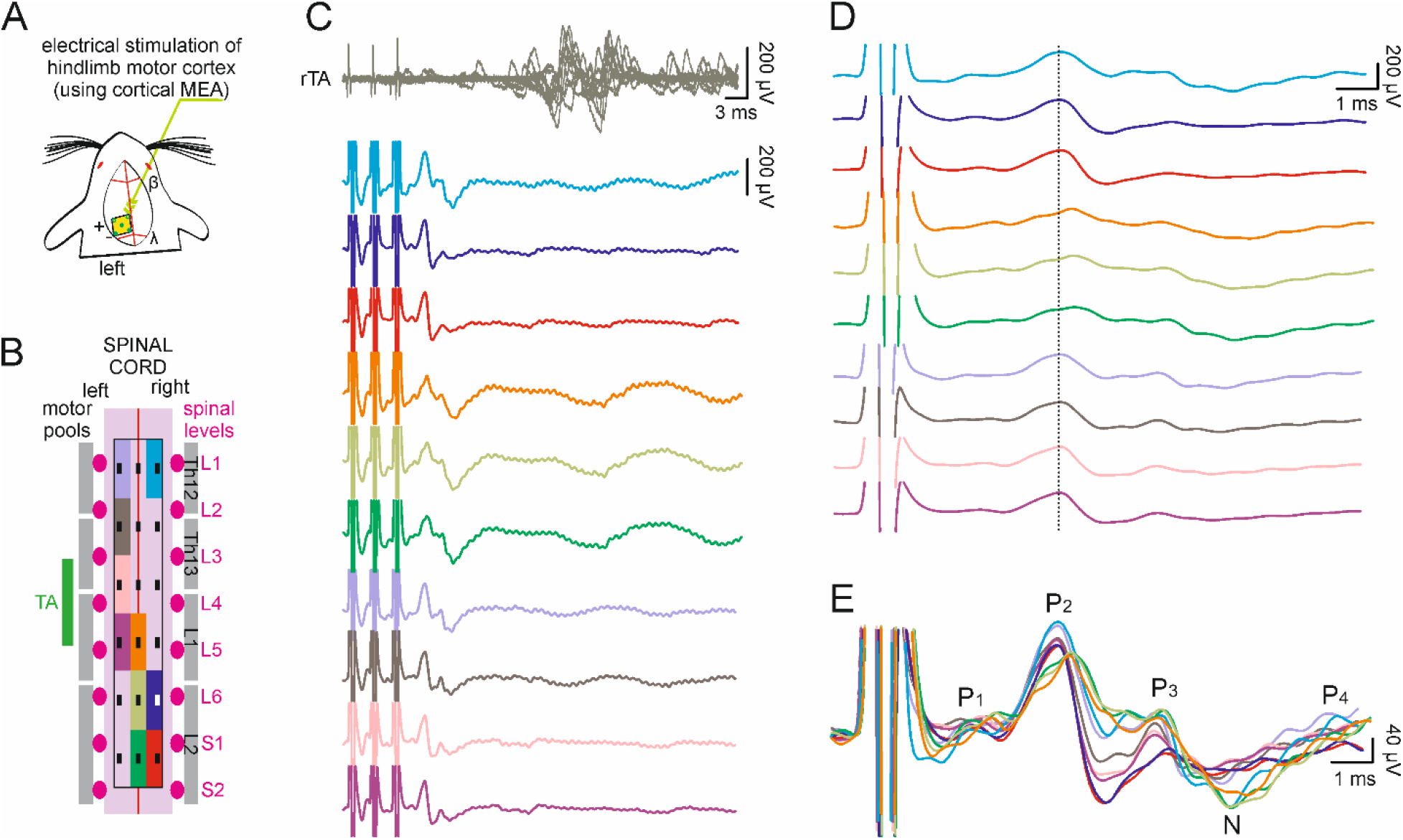
Selective electrical stimulation of the motor cortex evokes simultaneous CDPs, captured by multiple electrodes, along with delayed EMG responses from contralateral limb muscles. **A**. The cartoon schematizes the exposed brain with a calibrated drawing of the epidural array implanted over the left motor cortex under β and λ coordinates. **B**. A schematic view of our multielectrode array placed on the dorsal spinal cord. Dimensions of array and electrodes are calibrated based on the width and length of the cord, and the root entry positions. **C**. Simultaneous EMG recordings from TA of the contralateral hindlimb and CDPs, in response to a train of three biphasic pulses (333 Hz, intensity = 650 µA) delivered every 3 s to the left motor area. Simultaneous CDPs from 10 independent electrodes of the array are traced adopting the same color code as in B. Note that, as opposed to peripherally evoked CDPs, the ones elicited by supraspinal stimulation are conventionally represented with positive potentials pointing upwards. **D**. Cropped time-based scale of CDPs from C. The vertical dotted line connects latencies of the second negative wave for most CDPs. **E**. Superimposed traces follow the color code as in B, C and D, to show common volleys, which are indicated by the letters (P_1_, P_2_, P_3_, N, P_4_) conventionally adopted to describe CDPs elicited by supraspinal stimulation.

Along with the definition of waves in response to peripheral stimulation proposed by Bernhard (1952, 1953), the study will also adopt the terminology minted by Jordan and collaborators (Noga et al., 1995) to describe spinal potentials induced by descending brainstem stimulation as composed of positive peaks P_1_ to P_4_ and a negative slow wave, N. Time-to-peak values pooled from eight animals are P_1_ = 6.3 ± 3.6 ms; P_2_ = 9.6 ± 4.2 ms; P_3_ = 14.1 ± 3.9 ms; N = 18.0 ± 5.3 ms; P_4_ = 19.3 ± 4.5 ms.

Compared to the CDPs elicited by peripheral stimulation, cortically evoked CDPs occur entirely in the CNS. As a result, the peaks of presynaptic volleys are both positive (PV and P_1_), but the polarities of the postsynaptic components are swapped for peripherally-compared to supraspinally-evoked CDPs. Indeed, CDPs evoked by peripheral stimulation generate negative postsynaptic responses from spinal interneurons (Fig. 1A; N1-N2), whereas corticospinal pulses generate positive ones (Fig. 2E; P_2-4_). Contrariwise, the slower waves attributed to presynaptic inhibition of spinal neurons have a polarity that is positive for CDPs evoked by peripheral stimulation (P), but negative for the ones generated by corticospinal pulses (N). To keep a similar profile representation although the swapped polarities, peripherally evoked CDPs are conventionally reported with negative potentials pointing upwards, while the CDPs elicited by supraspinal stimulation are drawn with positive potentials pointing upwards (Noga et al., 1995).

### Dynamic stimulation selectively affects the P_2_ component of cortically induced CDPs

Once the possibility to record CDPs while selectively stimulating the motor area that controls hindlimbs was verified, we used the corticospinal platform to define whether the source of facilitation of motor responses due to DS was pre– or postsynaptic (Taccola et al., 2020a). In a sample experiment (Fig. 3A), CDPs were recorded from the dorsal spot (green) on the right side of the cord during a single electrical pulse (500 µA, pulse duration = 1 ms) delivered to the left cortical motor area every three seconds. During continuous stimulation of the motor cortex with test pulses (0.3 Hz), the four electrodes located at the extremities of the left and right columns of the epidural MEA were used to deliver DS to the whole lumbosacral enlargement, simultaneously supplying two EMG traces for 30 s on both sides of the array with rostro caudal opposite polarity and staggered onset (Taccola et al., 2020a). DS was supplied at sub-threshold intensity (300 µA, peak to peak) without any overt contraction of hindlimb muscles during stimulation. In Figure 3B, average CDPs of 30 sweeps, corresponding to pre-(black trace) and post-DS (red trace), are superimposed to show that DS did not affect the latency of each wave. Conversely, DS facilitated the amplitude of P_2_ by a 40 % (18.8 µV pre-DS vs. 26.3 µV post-DS), with no changes in P_1_ and P_3_ peaks amplitude.

**FIGURE 3.**
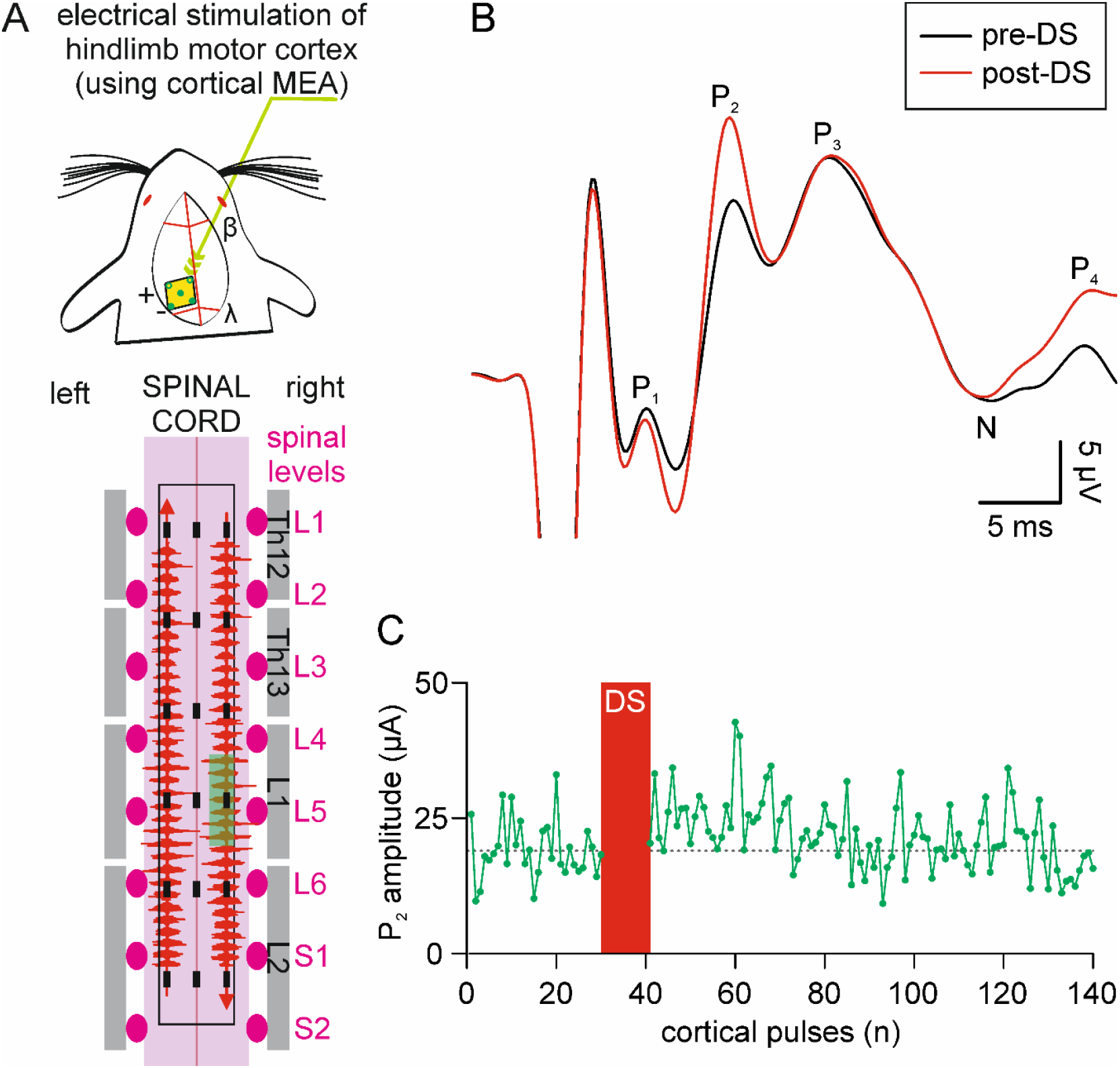
Dynamic stimulation of the dorsal cord selectively potentiates the P_2_ peak of cortically induced CDPs. **A**. Cortical (up) and spinal (down) multielectrode arrays composing the corticospinal platform adopted in the study. The highly varying protocol of stimulation named DS is applied at the extremities of the array, to both sides of the cord and with opposite polarity, as exemplified by the cartoon where vertical EMG traces in red are aligned to the two arrows on each side of the cord (arrowhead = anode). **B**. Superimposed averaged CDPs recorded from the right L5 spinal level, corresponding to the pale green field in the cartoon in A, and elicited by electrical stimulation of the left cortical motor area with single weak electrical pulses (500 µA, pulse duration = 1 ms) repeated every 3 seconds. Traces are averaged for 90 s, before (pre-DS; black trace) and after DS (post-DS; red trace). DS (300 µA, peak to peak) selectively increases the amplitude of P_2_, while P_1_ and P_3_ remain unaffected. **C**. For the same experiments as in B, the time course of P_2_ amplitude is plotted for 7 min of cortical stimulation. After control (30 sweeps, 90 s), DS was applied for 30 s (red rectangle), with artifacts of noisy stimulation covering all CDPs. Immediately after DS delivery, P_2_ amplitude transiently augments for about 30 sweeps (90 s) and then recovers to the average baseline value (dotted gray horizontal line centered around 19 µV).

In Fig. 3C, for the same experiment, we report the entire time course of the amplitude of cortically evoked P_2_ waves in control and after DS. During DS delivery, no CDPs could be acquired through the MEA due to some interference from the artifacts of stimulation. However, immediately after DS, the P_2_ peak remained higher for more than 90 s before recovering to mean pre-DS control values, here represented by the dotted gray line. The same observation was replicated in four animals with a significant P_2_ mean facilitation to 171 ± 45 % of pre-DS control (P = 0.043, paired t-test, n = 4), while P_1_ (P = 0.269, paired t-test, n = 4) and P_3_ (P= 0.198, paired t-test, n = 4) remained unaffected by DS delivery (625 ± 236 µA, peak to peak).

In summary, an innovative corticospinal platform revealed that DS selectively facilitates the peak of postsynaptic responses from those spinal interneurons that are the first to integrate corticospinal input, while maintains unchanged the extent of presynaptic input from corticospinal fibers, P_1_.

### P_2_ component of cortically induced CDPs appears in conjunction with the contraction of hindlimb muscles

We observed that DS selectively increased the amplitude of the P_2_ peak, maintaining this facilitation for up to three times the length of DS delivery. To ascertain the physiological meaning of the appearance of P_2_, input-output experiments were conducted during simultaneous CDP and EMG recordings. In Figure 4, a mean CDP was recorded from the midline at L5 spinal level (close to the motor pool for the TA, Fig, 1E), while simultaneously recording muscle activity in the contralateral TA. Trains of three bipolar pulses (frequency = 333 Hz; pulse duration = 0.2 ms) were applied at increasing intensities (400-550 µA) to the left cortical area for hindlimbs every 3 s.

**FIGURE 4.**
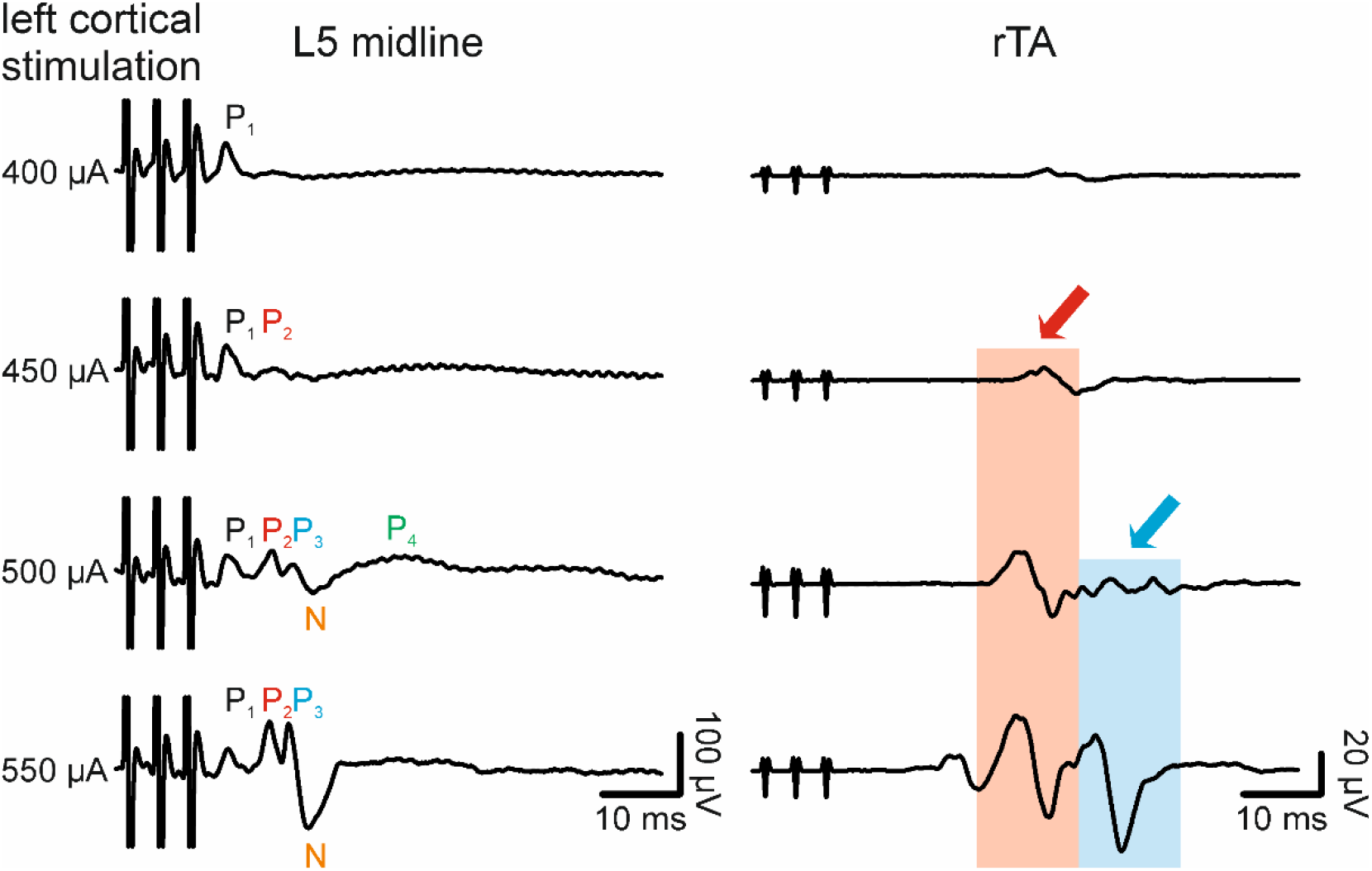
Increasing amplitude of cortical stimulation progressively recruit CDP components and full muscle response in the contralateral limb. Electrical stimulation of the left cortical area for hindlimbs (triplets at 333Hz every 3 s, pulse duration = 0.2 ms) at increasing amplitude (from 400 to 550 µA) simultaneously elicits average CDPs from the midline of L5 spinal level and EMG responses from the contralateral tibialis anterior muscle (TA, mean traces). Letters on the left identify the progressive appearance of CDP peaks, while colored arrows and pale fields on the right highlight the early and late EMG responses from right TA.

At the lowest intensity of stimulation, only the P_1_ peak is evoked from the cord. In correspondence to P_1_, a small deflection in the EMG baseline with a peak 28 ms after the artifact of stimulation, reflects a slight muscle recruitment in the contralateral leg (Fig. 4), indicating that, despite the presence of P_1_, descending input was unable to fully activate limb muscles. By increasing the intensity of cortical stimulation (450 µA), the P_2_ peak also appeared alongside a greater muscle recruitment (Fig. 4; pale red field). A further increase in the strength of stimulation (500 µA) produced a higher P_2_, which was accompanied by the appearance of waves P_3_ and P_4_ in the CDP volley, and a large N wave. The same pulse produced an additional and delayed EMG response with 36 ms of time to peak (Fig. 4; pale cyan field). The EMG signal was then maximally potentiated by the highest intensity of stimulation (550 µA), which also widened the amplitude of P_3_ and N. Notably, increasing intensities did not augment the amplitude of the P_1_ peak, signifying that the descending pre-synaptic volley was already saturated at intensities that elicited barely detectable EMG responses. The same observation was replicated in five rats, where the appearance of the P_2_ wave corresponds to the first slight deflection in the EMG baseline for a mean strength of stimulation of 634 ± 234 µA.

### Dynamic stimulation augments both EMG responses and the P_2_ component of cortically induced CDPs

DS has been reported to facilitate the motor output before and after a spinal injury (Taccola et al., 2020a, b; Culaclii et al., 2021). In the present study, DS increased the P_2_ component of cortically induced CDPs, which corresponds to the post-synaptic activation of spinal interneurons without any changes in the extent of the pre-synaptic descending input P_1_. On the other hand, we demonstrated that the appearance of the P_2_ peak parallels the slightest contraction of hindlimb muscles. To verify whether the potentiation of motor responses by DS mirrors an increase in the P_2_ component, simultaneous CDPs and EMGs responses were elicited by cortical pulses continuously applied before, during and after DS.

In an intact adult rat, weak cortical stimulation with short trains of three bipolar pulses (frequency = 333 Hz, pulse duration = 0.2 ms, amplitude = 780 µA) delivered to the left motor area elicited a small muscle contraction from the rTA, and a CDP volley, with a highest peak for P_2_, derived from the array electrode positioned on the left side of the cord, at the border between L5 and L6 spinal segments (Fig. 5 up-right). immediately after DS delivery (600 µA, peak to peak), the same cortical stimulation induced an increase to 267 % in the EMG response, which aligns with the augmentation in the amplitude of the P_2_ peak, which was 69 % of the pre-DS control (Fig. 5 up-left).

**FIGURE 5.**
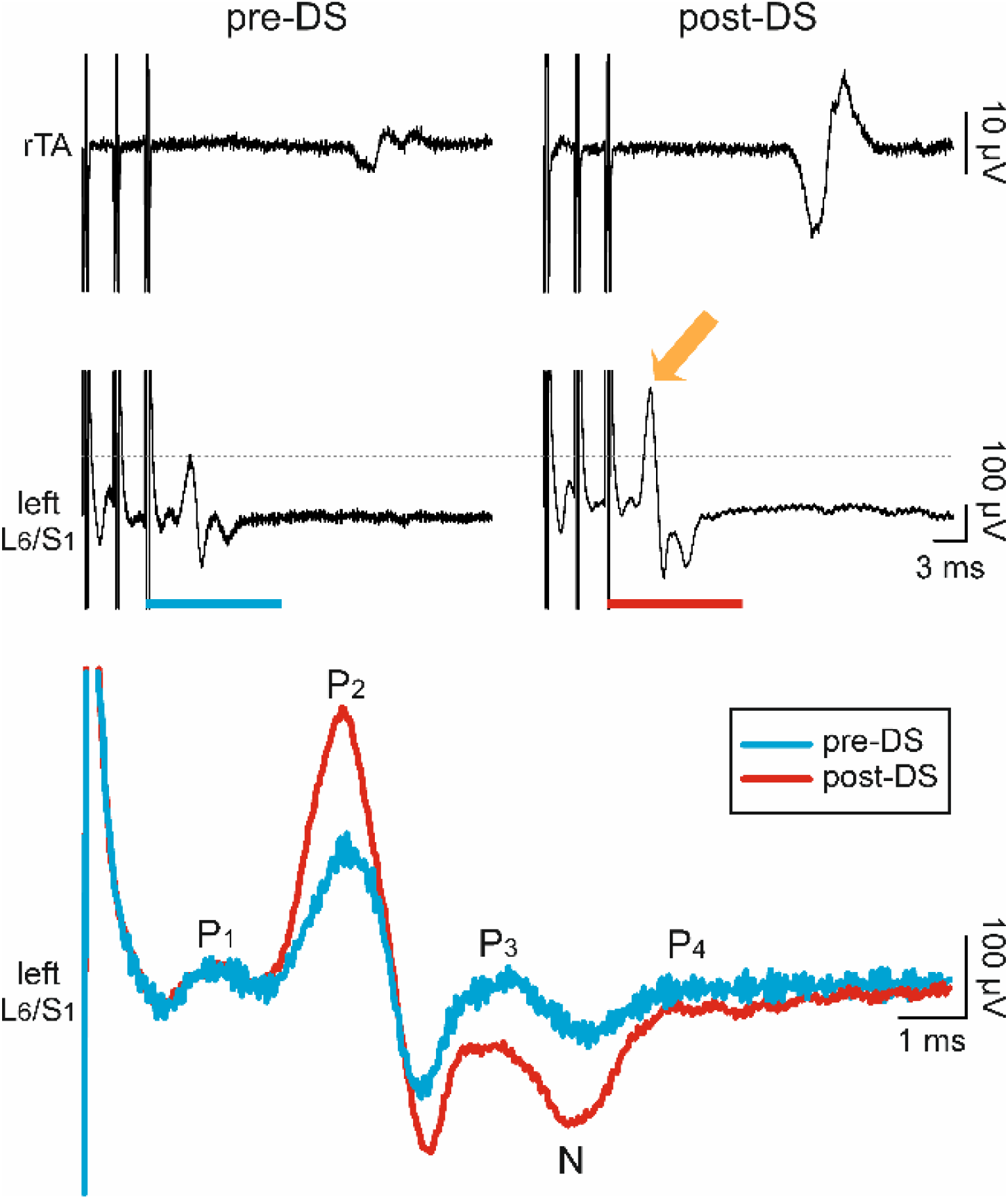
Dynamic stimulation facilitates cortically induced muscle responses and selectively increases the P_2_ peak of CDPs. Simultaneous mean EMG and CDPs traces are taken from right TA (top) and from the far-left side between L5 and L6 spinal segments (middle), in response to triplets (333 Hz, intensity = 780 µA, duration = 0.2 ms) of biphasic pulses delivered every 3 s to the left motor cortex. The mean CDP in control (left, pre-DS) is displayed with the amplitude of the P_2_ component indicated by the horizontal grey dotted line. Immediately after delivering DS to the cord (right, post-DS, DS intensity = 600 µV), motor responses are facilitated and the mean amplitude of the P_2_ peak is potentiated (orange arrow). Bottom, the averaged CDPs, corresponding to the trace segments indicated above by blue and red rectangles, are superimposed for control (blue trace, pre-DS) and right after DS (red trace, post-DS) and include conventional letters indicating peaks of the CDPs. Note that only the amplitude of the P_2_ component is almost twice the control, while other peaks remain unchanged.

Superimposed mean traces for 30 events before (blue trace) and immediately after (red trace) DS of the same experiment, displayed the selective potentiation of P_2_ after DS, yet unvarying amplitude of P_1_. Results from four rats indicated a mean DS facilitation of EMGs to 248 ± 167 %.

These results suggest that DS potentiates the post-synaptic P_2_ component of the CDP volley, which in turn corresponds to the facilitation of motor responses, as previously observed (Taccola et al., 2020a, b; Culaclii et al., 2021). Furthermore, DS increases the excitability of spinal interneurons without any direct effect on the magnitude of descending commands from the motor cortex.

## Discussion

### Epidural interfaces for recording and stimulating spinal networks

The multi electrode epidural interface adopted in this study has been developed and validated by a multidisciplinary team that provided a detailed engineering characterization (Gad et al., 2013; Chang et al., 2014) and tested efficacy in delivering epidural electric pulses to fully anesthetized intact (Taccola et al., 2020a) and SCI (Taccola et al., 2020a,b) animals, as well as in both, intact and chronically injured awake rats (Taccola et al., 2021). Platinum-titanium micro electrode array has been fabricated to ensure a low charge transfer resistance at the electrode-tissue interface (Chang et al., 2014). As a result, the high performance of stimulation is exemplified by the ability to supply complex protocols of stimulation with highly varying output (Taccola et al., 2022) that was simultaneously delivered at different sites of the interface. The latter feature was feasible due to the fully independent electrodes in the array, where each can be used to supply different pulses at the same time.

In addition to the versatility of stimulation, the low impedance feature of the electrodes of the interface led us to wonder whether we could also derive focal neural potentials from the surface of the dorsal cord. To challenge this hypothesis, we recurred to a classical neurophysiological sign of neuronal activity arising from the epidural cord, i.e. the CDPs elicited by peripheral stimulation.

The current study demonstrates that the ad-hoc designed multielectrode epidural interface is successful in simultaneously acquiring multisite neural signals, as peripherally evoked CDPs. Furthermore, the high-fidelity of recorded surface electric potentials allowed mapping of the rostrocaudal location of afferent terminals in the spinal cord, which correspond to the sites where the probability of finding synaptically coupled dorsal horn neurons is higher. Indeed, the magnitude of CDPs reflects the number of synaptic boutons in each afferent fiber (Koerber et al., 1990) and, also in our study, the highest CDPs evoked by stimulation of plantar and peroneal nerves appeared with a segmental location of afferent synapses, as previously identified by retrograde tracing experiments (Nicolopoulos-Stournaras and Iles., 1983; Koerber et al., 1990). In addition, the ability to detect the latency between afferent and presynaptic volleys is a touchstone for determining the intraspinal conduction of afferent input and, once the pulse conduction of fibers is also determined, providing a good estimate for the depth of afferent synapses located near the surface of the dorsal cord. Moreover, similar focal potentials from the cord were also recorded in response to corticospinal pulses.

In the current study, our multi electrode epidural interface was used for the first time to deliver highly varying patterns of stimulation to the four extremities of the epidural array, while simultaneously deriving cord potentials evoked by nerve or cortical stimulation before and after spinal neuromodulation with DS. To the best of our knowledge, only scattered evidence has been reported so far about recording multisite epidural potentials from the spinal cord (Parker et al., 2012; 2013), although their appearance did not resemble the classical multipeak profile obtained with ball probes or implanted microelectrodes (Brooks and Eccles, 1947; Bernhard, 1952, 1953; Eccles et al., 1954; Coombs et al., 1956). In addition, few studies reported the delivery of stimuli, as simple as square or rectangular pulses, through the same epidural interface used for recordings (Dietz et al., 2021). Indeed, pulse delivery and the acquisition of focal potentials from the surface of the spinal cord are challenged by several factors. First of all, the physiological curvature of the spinal cord requires optimal electrode compliance adopting arrays with a flexible frame, as the one in polyamide adopted in this study, to grant optimal adhesion to the surface. Secondly, the actual space between the dura mater and the spinal tissue is filled with conductive cerebrospinal fluid that spreads and attenuates any electrical signals. Third, the epidural array motion with back muscle contractions can lead to eventual misalignment of electrodes from original sites. Eventually, blood vessels on the surface of the spinal cord might represent scattered high-impedance spots interposed between the spinal cord and the epidural electrodes that reduce signal conduction from epidural electrodes to the spinal cord. In fact, in the current study, we lost signal from 8/9 electrodes at a time out of the 18 composing our interface, and these occasional failures sustain the need to adopt high-density electrodes when developing epidural arrays for the spinal cord.

Through the multi electrode array adopted in this study, we provided complex biomimetic patterns of stimulation and derived multi-site classical CDPs of different amplitude in relation to the electrode position away from the direct synaptic source. This observation can drive further technological advancements, leading to future clinical interfaces that can provide a real-time spinal read-out of the efficiency of the pulses supplied.

### Insights on the mechanisms of spinal neuromodulation

Although the mechanisms of spinal electrical neuromodulation are still elusive, several scientists repute that electrical stimulation of the spinal cord through epidural dorsal electrodes is a mere activation of Aβ-fibers in dorsal roots (DRs). Albeit the activation of Aβ-fibers is inevitable, since they are low-threshold afferents characterized by a large diameter, high myelination and ability for rapid signal conduction (Holsheimer, 2002), we support the view that spinal neuromodulation is an infinitely more complex phenomenon, including an extensive neuronal network infrastructure, anatomically and physiologically (i.e. neurons, axons, glia and extracellular fluid (Taccola et al., 2018). This concept has been clearly demonstrated, among other studies, by the pioneering work of Swiontek and collaborators (1976) showing that electrical pulses applied through a clinical stimulator placed on the dorsal surface of spinal cords explanted from human cadavers, evoked current density patterns detectable across the entire cord and even in the inner gray matter. Mehta and colleagues (Moore et al., 2017) recorded from cortical neuronal dendrites, demonstrating sub– and supra-threshold dendritic membrane potential from different segments of dendrites in unanesthetized freely behaving rats with firing rates several-fold larger than action potentials generated by the soma. In particular, subthreshold dendritic membrane potentials fluctuations that were substantially larger than dendritic action potentials could profoundly alter synaptic plasticity, altering the nature of neural coding and memory. Although these varying membrane responses unique to different segments of the dendritic branches were observed in cortical neurons have not been reported spinal neurons, it suggest caution in assuming that the more critical factors in controlling movement and the mechanisms of neuroplasticty is likely attributable to the size of an axon and the fundamental building block of neural information processing. These results have important implications for neural coding and plasticity. Furthermore, in the present study, through the same array used for spinal stimulation and placed on the surface of the dorsal midline, we derived electrical potentials of roughly 120-230 µV (Table 1-5, aN2) that arose from action potentials (70 – 80 mV; Husch et al., 2015) generated in second and third order spinal interneurons (N2, P_3_, P_4_ waves). These values reflect an attenuation of 300 – 600 times the original signal, as it crosses the spinal tissue before reaching the dorsal surface. As a result, stimulating intensities above 6 V, in the range of the ones routinely delivered in preclinical studies (Courtine et al., 2009) should be sufficient to directly depolarize second and third order interneurons of around 20 – 30 mV, hence triggering action potentials (Husch et al., 2015) regardless the presence of incoming afferent input through dorsal roots. Acknowledging that spinal neuromodulation is much more than a mere stimulation of afferent fibers suggests why functional recoveries obtained with an epidural array (Harkema et al., 2011) have not been replicated so far through the selective stimulation of dorsal roots, dorsal root ganglia or even peripheral nerves.

### Corticospinal input is facilitated by direct stimulation of its spinal targets

The present study demonstrates that spinal interneurons that integrate descending input from the motor cortex are the first targets of neuromodulation with noisy patterns of stimulation, which facilitate cortico-spinal excitability and motor responses. Although all of the mechanisms of spinal neuromodulation remain unknown, our contribution demonstrates that the broad excitation of the dorsal cord due to neuromodulation acts on a group of spinal interneurons responsible for the P_2_ wave, by bringing them to threshold to fully activate more motor units in a given motor pool in response to otherwise inadequate cortico-spinal input. These novel results add up to the facilitation of volitional commands for motor control induced by spinal neuromodulation in neurological disturbances characterized, at least in part, by the impairment of cortico-spinal tracts.

In the current study, the classical neurophysiological approach implemented through a specifically designed corticospinal MEA platform, enables the dissection of the pre– and post-synaptic action sites of the motor output. The pre-synaptic volley of afferent input elicited by peripheral nerve stimulation corresponds to the short latency (around 2 ms) positive peak, with an amplitude that increased during delivery of stronger pulses, reminiscent of the recruitment curve of H responses (Knikou, 2008). Contrariwise, in the present study, the descending pre-synaptic volley elicited by cortical stimulation was already saturated at lower intensities, barely able to elicit detectable EMG responses, and did not change even with stronger pulses. The present finding might suggest that to fully titrate the amplitude of presynaptic input in response to cortical stimulation, then is required a cortical stimulation protocol of lower intensity than that used here. On the other hand, despite the unchanged amplitude of descending input, increasing strengths of cortical pulses might recruit additional corticospinal collaterals impinging onto the spinal interneurons responsible for the generation of the P_2_ wave and linked to motor output facilitation.

Beside their functional role, the current study did not identify the interneurons responsible for the P_2_ wave. The latency (around 9 ms) of activation by cortical input suggests that this interneuronal pool has a direct, oligo-synaptic connection with supraspinal centers for motor control. Noteworthy, a distinct glutamatergic immediate executor module (iEM) that transforms descending commands into rhythmic patterns for locomotion has been recently identified in the spinal cord (Hsu et al., 2023). The location of both, iEM and the functional pool that generates the P_2_ wave in our study, parallels the location of groups of spinal interneurons that generate a descending volley on the surface of the middle aspect of the dorsal lumbar column in response to MLR stimulation (Noga et al., 1995).

On the other hand, it seems unlikely that spinal neuromodulation can strengthen axon conduction through fibers that are non-electrically competent to perceive weak descending input. Indeed, the present data demonstrate that the pre-synaptic volley peaking at P_1_, remains unmodified even by increasing intensities of DS, regardless of the progressive muscle recruitment due to higher strengths of stimulation. This suggests that the effect of DS in facilitating cortico-spinal input does not involve any potentiation of supraspinal volleys generating P_1_.

On the contrary, the P_2_ wave appeared in correspondence to an adequate EMG response, and the P_2_ peak amplitude increased along with stronger bouts of cortical stimulation that eventually elicited a greater motor output, which also included a rise in the P_3_ peak linked to the appearance of a late EMG response. Interestingly, without any changes in the extent of P_3_, DS selectively potentiates both, the post-synaptic P_2_ component of the CDP volley and the EMG responses, as previously observed (Taccola et al., 2020a, b; Culaclii et al., 2021).

In accordance with this view, recovery of volitional motor control after SCI has been obtained by the direct neuromodulation of sublesional lumbar networks that contain the post-synaptic targets of descending commands (Harkema et al., 2011; Gerasimenko et al., 2015; Angeli et al., 2018; Gill et al., 2018; Wagner et al., 2018).

### Spinal cord stimulation for supraspinal motor disorders

A hot topic in motor neurorehabilitation is the different strategies to stimulate the spinal cord to facilitate recovery of different motor and autonomic functions. This intervention was initially targeted to the spinal cord following spinal injury (Harkema et al., 2011; Gerasimenko et al 2015; Sayenko et al., 2022) to reactivate and train spinal locomotor circuits disconnected from brain commands (Taccola et al., 2018). Surprisingly, gait improvements have been recently reported in neuromotor disturbances secondary to brain dysfunctions, as in advanced Parkinson’s disease (PD; Pinto de Souza et al., 2017; Samotus et al., 2018), restless legs syndrome (De Vloo et al., 2019), primary orthostatic tremor (Lamy et al., 2021), multiple system atrophy (Millar Vernetti, 2022) and stroke (Awosika et al., 2020).

Symptomatic effects of electrical stimulation in subjects with PD have been mainly ascribed to the modulation of ascending sensory pathways, which increase cortical and thalamic output (Thiriez et al., 2014). In addition to this view, data presented in the current study suggests that the increased recruitment of spinal networks due to spinal neuromodulation might mitigate motor abnormalities that follow an impaired corticospinal input, a non-rare manifestation in PD (Xu et al., 2020).

In summary, the present study casts some light on the mechanisms of electrical stimulation of the cord in enhancing the recruitment of spinal networks, thus supporting the use of neurostimulation whenever the motor output is jeopardized by a weak volitional input, due to a partial disconnection from supraspinal structures and/or neuronal brain dysfunctions.

## Acknowledgments

GT is also grateful to Dr. Philippa Warren for her assistance with animal experiments and analysis. GT owes Dr. Elisa Ius for her excellent assistance in preparing the manuscript.

## Conflict of interest

VRE, a researcher on the study team holds shareholder interest in Onward and holds certain inventorship rights on intellectual property licensed by The Regents of the University of California to Onward. VRE, and PG, researchers on the study team hold shareholder interest in SpineX. WL holds shareholder interest in Aneuvo as well as inventorship of IPs licensed by The Regents of the University of California to Aneuvo.

## Funding

GT is supported by funding from the European Union’s Horizon 2020 Research and Innovation Program under the Marie Sklodowska-Curie (grant agreement No. 661452). This research was also funded in part by NIH/NIBIB U01EB007615, NIH/NINDS 1U01NS113871-01, International Spinal Research Trust (STR104), Walkabout Foundation, Dana & Albert R. Broccoli Charitable Foundation and Nanette and Burt Forester, including matching by PwC LLP and Roberta Wilson.

